# Methods for Running a Successful Women-in-STEM Organization on an Academic Campus

**DOI:** 10.1101/2020.02.20.958629

**Authors:** Deborah D. Rupert, Alexandra C. Nowlan, Oliver H. Tam, Molly Gale Hammell

## Abstract

The current academic culture facing women in science, technology, engineering, and math (STEM) fields in the United States has sparked the formation of grassroots advocacy groups to empower female scientists-in-training. However, the impact of these initiatives often goes unmeasured and underappreciated. Our Women in Science and Engineering (WiSE) organization serves post-doctoral researchers, graduate students, and research technicians (trainees) at a private research institute for biological sciences. Here we propose the following guidelines for cultivating a successful women-in-STEM-focused group based upon survey results from our own scientific community as well as the experience of our WiSE group leaders. We hope these recommendations can provide guidance to advocacy groups at other research and academic organizations that wish to strengthen their efforts. While our own group specifically focuses on the underrepresented state of women in science, we hope these guidelines may be adapted and applied to groups that advocate for any minority group within the greater scientific community (i.e. those of gender, race/ethnicity, socioeconomic background, sexual orientation, etc.).

## Introduction

Substantial data show that while the number of undergraduate and graduate degrees in STEM awarded to women is roughly equal to the number awarded to men, women remain underrepresented in professional leadership positions both in academia and industry.^1–3^ Compared to men, women are more likely to be targets of hiring bias, micro-aggressions, and sexual harassment, and receive fewer invitations to publish and present their research.^3–5^ These factors have direct consequences on career outcomes and long-term retention of women in STEM fields.^6^ Indeed, not just overt bias, but ambivalence towards sexism and bias has been reported to negatively affect female trainees.^7^ The marginalization of women in STEM was publicly recognized in a landmark study conducted at MIT in the late 1990s.^8^ Many of the identified inequities have persisted, leaving female scientists dissatisfied with the limited extent of reform within academic institutions and STEM communities more generally. Moreover, these inequities negatively affect the scientific community at large; driving female talent out of science restricts scientific progress and has larger consequences for the health of the general population when medically-relevant research at both the bench and public health levels are gender-restrictive.^9^

In recent years, there has been considerable investment in initiatives to support women’s advancement in STEM and spark change from the bottom up.^10^ These grassroots organizations are critical for empowering female scientists-in-training. At Cold Spring Harbor Laboratory (CSHL), a private institute for biological research and education, our Women in Science and Engineering (WiSE) organization is one such group created by and for trainees: post-doctoral researchers, graduate students, and research technicians. Our aim is to foster a more supportive, collaborative and equal-minded scientific community by providing a platform for professional development, education, and empowerment.

Our recommendations are founded in over 4 years of experience establishing a women-in-STEM advocacy group on an academic campus. However, to further substantiate our views, we chose to conduct a quality assurance survey of our campus to determine the extent to which our own group had been successful in meeting its goal and the current community attitude towards our initiative. According to anonymous community feedback, approximately 80% of campus-wide, survey participants considered CSHL’s WiSE program to be moderately or very successful based on their understanding of our organizations’ mission and goals.

Examining this feedback has allowed us to evaluate our group’s accomplishments and short-comings. We use the conclusions about these strengths and weakness as well as our own experience to propose the following suggestions for women-in-STEM advocacy groups that are starting up or who wish to bolster their own efforts.

## Methods

### Survey Distribution

Our survey was distributed to all members of the campus and probed participants’ understanding of the goals, the relative success in meeting those goals, and the perceived weaknesses and strengths of our group. In addition, we gathered demographic characteristics of our respondents (Fig 1) in order to understand what factors might affect those perceptions. The majority of survey participants did not consider themselves active WiSE members (73.81% were non-members, 25% members, and 1.19% former members), which is helpful for getting feedback from the greater community in which our group works i.e. perspectives external to the WiSE group but within the CSHL community.

**Figure 1.**
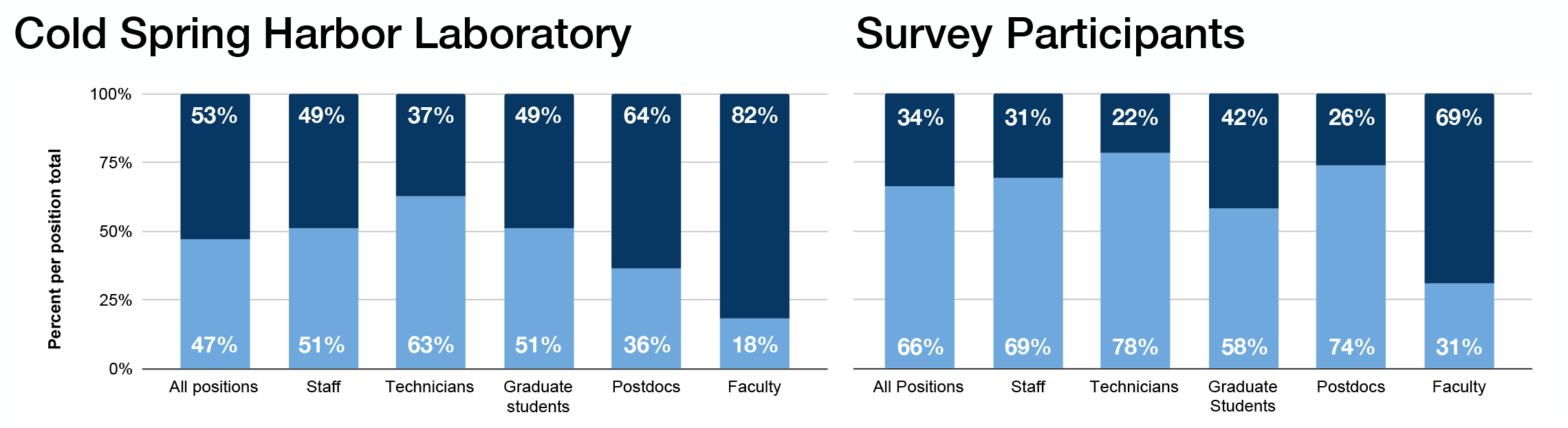
Demographics of greater CSHL community (2018) alongside survey participants. As seen here, gender parity among graduate students on the CSHL campus reflects those across the nation. Similarly, lack of parity among support staff positions (more female than male), postdoctoral researchers, and faculty (more male than female) also fits with national trends and the ‘leaky pipeline’ theory. In comparison, participants in our 2018 survey were more likely to be female.

Our survey was entirely anonymous and voluntary. Responses were collected using a free, online platform (SurveyMonkey) over the course of a month’s time. A total of 167 individuals participated and the responses of all but two individuals were included in our analysis; those excluded answered less than half of the questions making up the survey. Where participants chose not to answer questions, they were excluded from those sections of the analysis as necessary. As a first step, we classified the positions of all survey respondents (Fig 1), which mainly fell into the categories, of: faculty, staff, trainees, and others. The data was then manually examined and cleaned in order to properly classify individuals that responded with “other”; the most common instance of this was staff members providing more specific position information. For example, manual cleaning for “what is your current position (at CSHL)?” might include a response such as “other: public relations” which would be re-categorized as “staff”.

### Statistical Analyses

All statistical comparisons of male to female responders within a single category were calculated using Fisher’s Exact Test. When relative participation rates across categories were compared, a paired T-test was performed. A Benjamini-Hochberg correction was used to account for multiple hypothesis testing, such that reported probabilities represent FDR-corrected Q-values.

## Results

### Define and communicate the goal(s) of the group

Assessment of an organization’s success is best measured against a clear mission statement. We suggest formalizing both broad and event-specific goals in writing and actively revisiting these pieces of writing to adapt them as the group evolves i.e. annually when new leadership is elected. For example, our group’s general mission statement is:

> “To build a more supportive, collaborative, and equal scientific community for all. We provide a platform for professional development and empowerment through mentorship, career planning, and educational opportunities tailored toward issues disproportionately affecting women.”

While the ‘mission’ for a specific event might read as:

> “The WiSE Retreat is an annual, day-long meeting that welcomes scientists-in-training at CSHL to attend lectures on gender disparity. The goals of the day are to: 1. familiarize the WiSE group and our greater community with gender disparity literature; 2. encourage the application of scientific scrutiny to this body of literature; 3. stimulate continued self-education on this topic.”

Our group broadcasts it’s goals and announces WiSE-hosted, on- and off- campus events through a website platform (*www.cshlwise.org*), Twitter account (@CSHLWISE), Instagram (@WISE_CSHL), Facebook (/WISECSHL), and campus-specific Slack account. Advocacy groups (i.e. institutional initiatives or student run groups that aim to promote equity for scientists of minority status) should be cognizant that their broadcasts will suffer from self-selection bias on the part of followers; thus, some subpopulations of the academic campus will be better informed than others.

Although we strive to effectively communicate the inclusive nature of our WiSE group and our mission to foster an equal, supportive, and collaborative scientific environment, we have found that doing so is more readily said than done.

In order to understand whether we had effectively communicated our mission to the campus, we asked survey participants to select the best fit to their interpretation of WiSE’s goals from three potential descriptions. The three options stated: 1) “WiSE is an open group for everyone at CSHL,” 2) “WiSE is a female-only group”, and 3) “WiSE is a political group.” While WiSE promotes inclusion of all members of the CSHL community to the WiSE group, we found the group has not fully succeeded in conveying this message. As demonstrated in our survey data, despite our vigorous attempts on social media platforms to promote the message that WiSE is an open group for all CSHL members, over 34% of our participants incorrectly stated the aim of our group, choosing options 2 or 3 above. As demonstrated by our own survey data, over 34% of our participants incorrectly stated the aim of our group. Moreover, male community members were more likely overall to state that WiSE was only open to women (P<0.006, Fig 2). We recognize this to be, in part, a shortcoming of our own efforts to communicate and promote the inclusion of all campus members. We encourage fellow organizations to evaluate their communication strategies, examine when they fall short, and determine how to address audiences that may be less familiar with the group.

**Figure 2.**
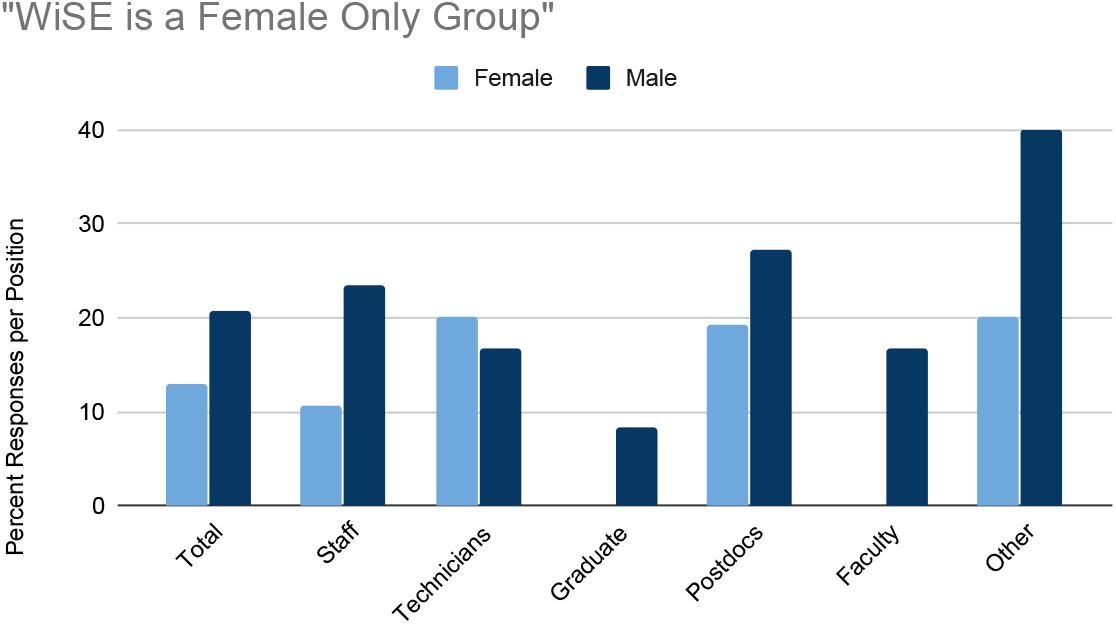
Perception of group’s mission by gendered position. Across all members of the academic community, males were more likely to believe WiSE to be a female-only group (Paired T-test P<0.006).

### Carefully structure the organization

Our WiSE group addresses the present needs of the CSHL community through five subcommittees: Institutional Initiatives, Professional Development, In-House Education, Mental Health, and Outreach. Each committee reaches toward an individual goal (Table 1), while all five work together to support female researchers on our campus in reaching the height of their potential.

**Table 1.** List of events hosted by our group to date, broken down by category. Events reflect the pillars of our organization: institutional initiatives, professional development, outreach, education, and mental health. To learn more about these events please visit our website: cshlwise.org

| Branch of Our Organization | Goal | Example Events |
| --- | --- | --- |
| **Institutional Initiatives** | Highlight the accomplishments of prominent female scientists in biology and provide networking opportunities for our trainees | McClintock campus wide lectures |
|  |  | Women in Biology Speakers List |
|  |  | WiSE Mentorship awards |
|  |  | Breakfast with invited female speakers |
| **Professional Development** | Promote the professional growth of our trainees | Negotiation workshop |
|  |  | Financial management workshop |
|  |  | Presentation skills workshops |
|  |  | Practice talks for graduate students and postdocs |
| **Outreach** | Grow the next generation of scientists, feminists, and their supporters committed to equality in STEM | Girl Scouts Brain Awareness Week |
|  |  | Basic coding bootcamp |
|  |  | Summer camp |
|  |  | Wiki-edit-a-thon |
|  |  | Science cafes in high schools |
| **Education** | Systematically review data-driven literature examining gender disparities in STEM | Journal clubs |
|  |  | Annual education retreat |
| **Mental Health** | Combat the mental health crisis endured by trainees, which disproportionately affects women | Open Mic mental health night |

We encourage our counterparts at other academic institutions to carefully consider their primary goals and how to best go about accomplishing these given the limitations of their women-power, resources, and institutional support. Our own group was originally established with 3 of its now 5 subcommittees which have been modified as the group’s leadership evolves.

We further recommend careful consideration of how the structure of the organization can best benefit its members. Leaders should have realistic conversations about board member expectations when inviting new members to take on leadership roles. For example, we ask board members to commit to a position for 1-2 years, whereas certain roles (i.e. president) can only be held by a board member with prior experience in a different position. At the same time, being able to adapt board roles to fit the time-commitments and strengths of future leaders is critical for achieving realistic progress. It is important to remember that our members are scientists first and foremost, and ideally their level of involvement in advocacy should be balanced such that it does not interfere with their research progress.

Our group’s leadership is structured to promote gradual increases in responsibilities (Fig 3). Leaders develop and practice critical skills such as project management, communication, and professionalism through their work in WiSE. When senior leadership members transition to the next stage of their training, their roles and responsibilities must be handed off to the next leader with clear expectations and an open-door policy. Mentoring within the group is essential so that senior leaders can pass the torch on to younger members. Doing so benefits both parties, allowing younger members to acquire more responsibility and allowing senior members to release that responsibility when it becomes too demanding in conjunction with their career demands. Volunteer advocacy gives scientific trainees real life experience with leadership, time management, and negotiation, all of which are integral to a career in science.^11^ Thus we argue that grassroots organizations like our own are a fundamental part of trainee development in addition to the scientific development they undergo.

**Figure 3.**
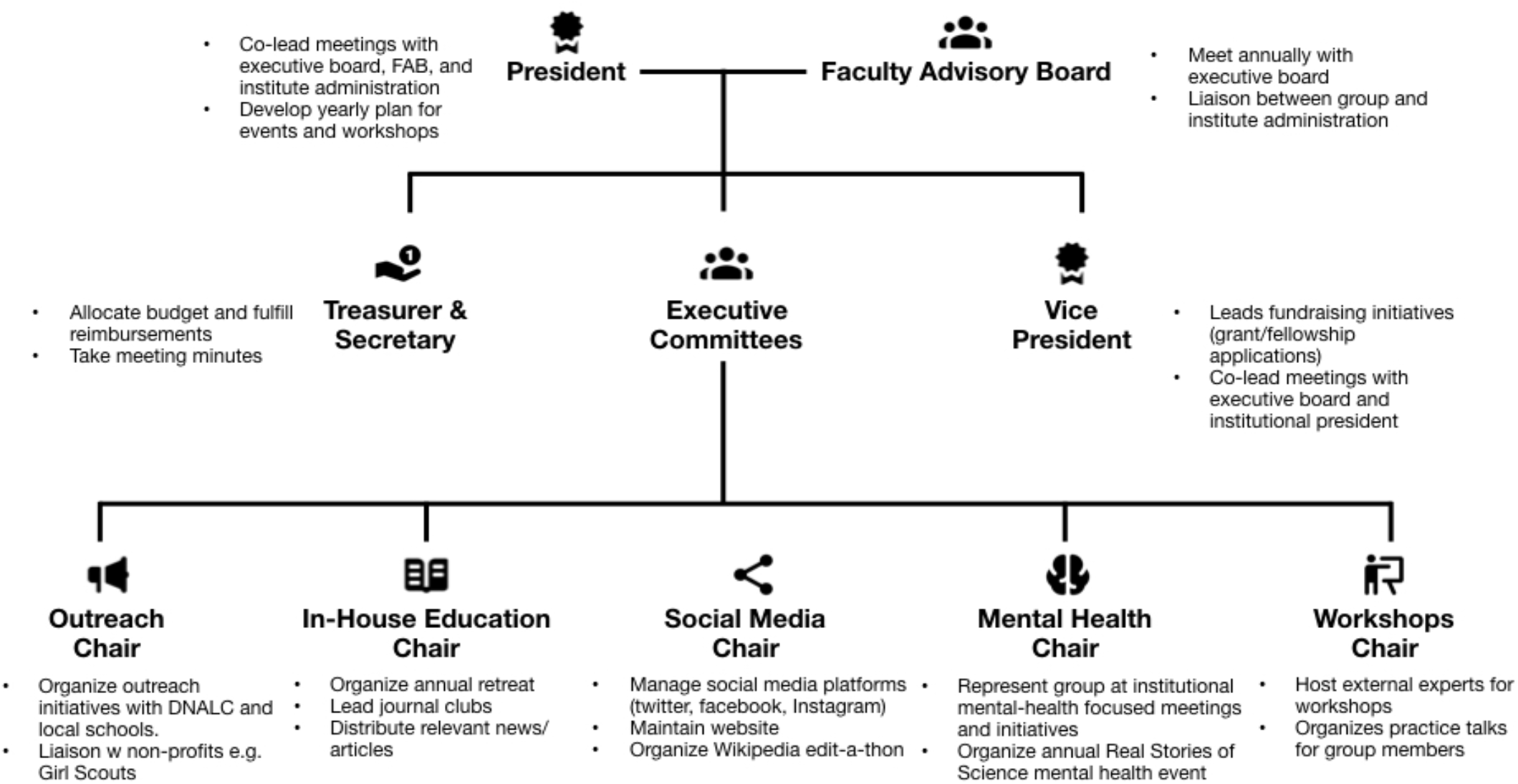
Schematic of WiSE organizational leadership. WiSE is a trainee led group with students, postdoctoral fellows, and technicians filling all of the leadership roles outside of the Faculty Advisory Board. WiSE members are encouraged to volunteer and participate in the events organized by the executive committee chairs which oversee the subcommittees/branches detailed above. Active members typically go on to assume a role on the board; this allows for many leadership opportunities for students and ensures that the group remains active as responsibilities are handed off to subsequent group leaders.

### Harness the strengths of members

Our group is composed of women, and allies of women in STEM, at various levels of scientific training (technicians, graduate students, and postdocs) and across various fields of biological sciences (cancer biology, neuroscience, genomics, etc.). We see strength in our ability to recognize members as having unique backgrounds, interests, and skill-sets (scientific and otherwise) and harness this diversity. This is reflected in our hosted events (Table 1). For example, members with expertise in computer programming have led hands-on, basic coding bootcamps for an audience of middle-school-aged, young women. A member with a proclivity for social media organized a “How To” wiki-edit-a-thon to help expand digital media articles on prominent female scientists. Members with an interest in teaching careers have applied and expanded upon their prior experience through our educational outreach events. The various interests of our members are also reflected in the organization of our subcommittees. For example, our in-house education committee was added to distinguish external outreach events from our efforts to educate our own community. The addition of a committee is a large undertaking that must be carefully considered by the board and backed by an appropriate level of commitment by the founding members. Similarly, when the group’s efforts become spread too thin, removal of specific events or committees must be considered to maintain the integrity of the group’s efforts.

We encourage advocacy groups to balance the specific interests of active members with the assurance that these events meet the expectations of the target audience. At the same time, recognizing that targeted audience subpopulations vary by event is fundamental for allowing specialization of one’s initiatives and breadth of the organization’s reach as a whole.

### Identify goals common to institutions and groups

The degree to which an institute recognizes or resists gender disparity issues undoubtedly varies. Despite this, common objectives between women in STEM-focused groups and their institutions can help benefit both parties. For example, institutions are self-motivated to attract strong scientific minds and “big name” guest lecturers. WiSE groups should be mindful of the goals of their underlying institution, the decision-making leadership and stakeholders within that institution, and the strength of other groups that it chooses to collaborate with. Impact can best be achieved where all parties gain. For example, our WiSE organization has negotiated for institutional support (both financial and logistic) to host two prominent female scientists to give guest lectures through the McClintock Lecture series, in which we honor the legacy of Nobel Laureate and CSHL scientist Barbara McClintock (see Table 1 and Fig 6).

We hypothesized, and confirmed, that participants with academic positions would be more likely to attend the McClintock lecture series (academic events) over other WiSE events (Fig 4). We found both male and female participants across positions do show a preference for academic events (Fisher’s Exact, P<0.008). Understanding that academically interesting events can be used as a draw to pull in a wider audience can be a powerful way to share the message of women in STEM groups beyond those who would regularly be exposed to it. As such, the McClintock lectures, which draw widely from the CSHL community, provide a platform for promoting WiSE and announcing other upcoming events.

**Figure 4.**
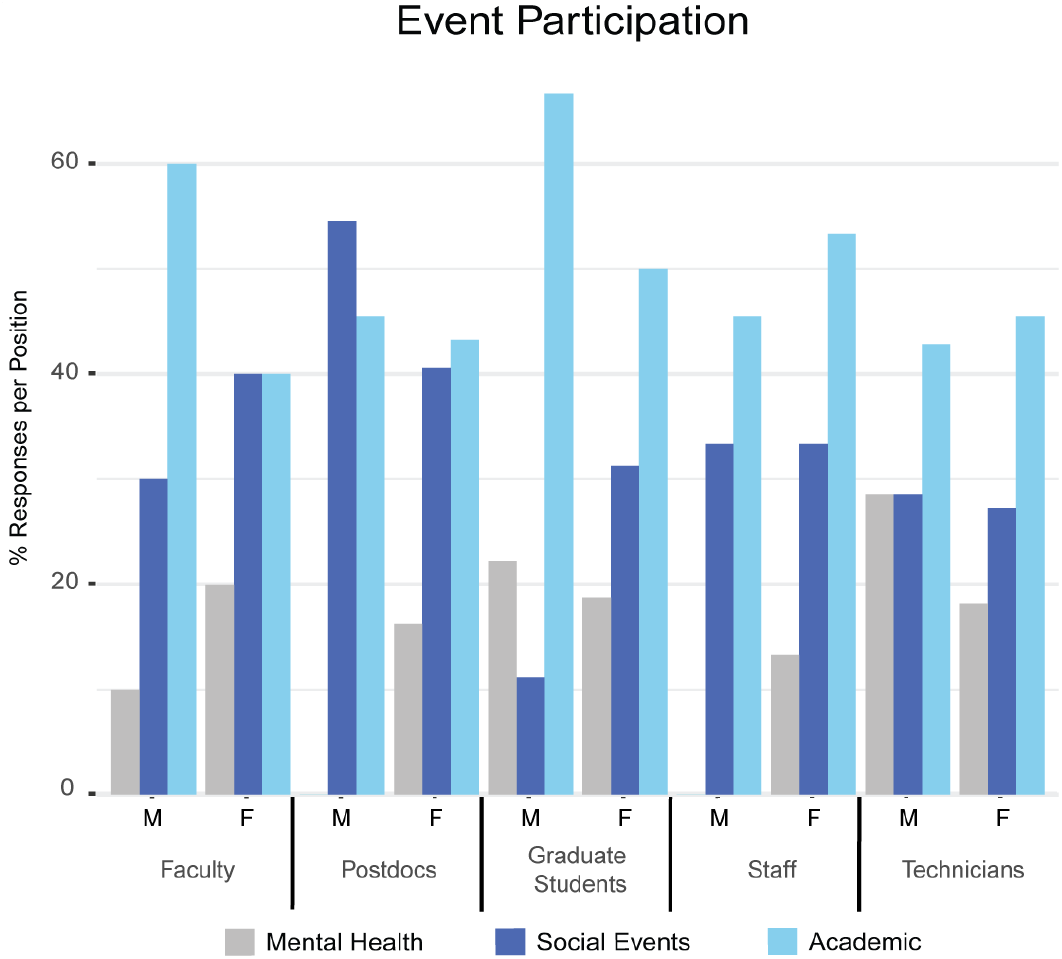
Event participation rates by gendered position. Response rates were normalized to the total number of responses for a given gendered position across all event types. Event types were pooled into categories of “mental health”, “social events”, and “academic”. Female participants were more likely to participate overall (T-test P<0.028) although the academic events showed even representation across gender. Male survey participants were least likely to attend mental health events (P<0.042, Fisher’s Exact)

We additionally found that graduate students were more likely to attend mental health events (Fisher’s Exact, P<0.014), while postdoctoral fellows were more likely to attend social events (Fisher’s Exact, P<0.014). This suggests that both events are important to different members of our community.

Lastly, we highlight the value that advocacy groups add to their institutions. Trainees expect this type of community support on campus, and look to the leadership to endorse this overdue shift in academic culture towards supportive and inclusive environments.^1,2,10^ Members of our group are frequently called upon by the institute in matters of public relations to represent our goals as well as those of the larger institution. Accepting these opportunities whenever possible is paramount to cultivating a productive relationship with the administration.

### Promote diversity within the group

We believe that attracting diversity to one’s group in terms of gender, position, and racial/ethnic background is critical for the success of the organization.

When we investigated the factors that might discourage membership, we lumped response options into the following categories: know how (including “I’ve thought about getting involved but don’t know how” and “I don’t think I could be helpful for the group”), negative opinions (including, “I don’t think the group is productive”, “I think the group is too political in nature”, and “I don’t think it is necessary to have a women-focused group on campus”), lack of time, and lack of applicability.

Across academic positions (faculty, post-docs, technicians, and staff) men were more likely to believe the WiSE group was not for them, as compared to women (P<0.14, trend, Fisher’s Exact). Men were also more likely to indicate that they did not know how to get involved (P<0.18, trend, Fisher’s Exact). While both results were not statistically significant, they persisted across positions (Fig 5). Interestingly, male and female participants across positions are equally likely to have a negative opinion of the group influencing their decision to not participate, although this was a small percentage of all respondents.

**Figure 5.**
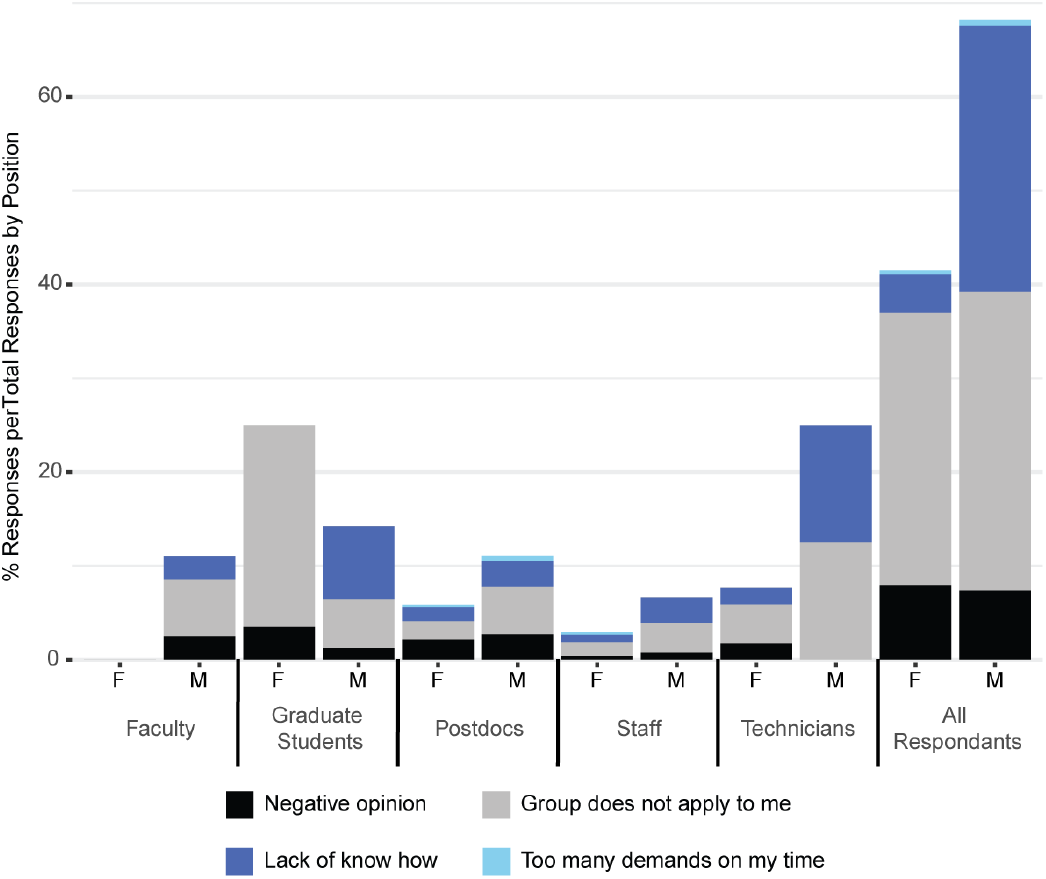
Response rates for reasons given for lack of involvement. Total responses for each gendered position were normalized to the total number of response reasons given across options for that gendered position and the total number of gendered participants in the survey. Men were more likely to believe the WiSE group was not “for” them (P<0.14, trend, Fisher’s Exact) and were more likely to indicate that they did not know how to get involved (P<0.18, trend, Fisher’s Exact).

We note that participants that identified as non-member female graduate students were the most likely compared to other positions to feel that the group did not apply to them (Fig 5). Since female graduate students formed a relatively large fraction of the overall survey participants, they also formed most of the respondents listed as “all” in the last column. We cannot determine, but suspect, that these non-member female graduate students do not feel WiSE applies to them because the perceived values of the WiSE group do not match their own, e.g. they do not self-identify as feminists.

Our data is privy to self-selection, i.e. we expect to get fewer responses from those who hold a negative opinion of the group or who do not feel the group applies to them. This is evident in the relative response rates of academic track males and female participants compared to the size of these populations on the CSHL campus (see Fig 1).

However, we conclude from these findings that WiSE groups, our own included, often develop tunnel vision and fail to engage untapped sources of support. For example, staff personnel on our campus pointed out the lack of roles for them to play in the group (“Lack of know how”, Fig 5); this is a large body of supporters whose assistance remains underappreciated and untapped. Male allies were also more likely to state that they did not to have a clear understanding of how to get involved or provide support (Fig 5).

These data are paralleled by a lack of event participation by male campus members; when participation was broken down into male and female responders, we found that male faculty, graduate students, and staff were less likely across the board to participate in all event types than their female counterparts (P<0.028, Paired T-Test, Fig 4). Academic lectures represented the one event with near equal participation, at an average of ~36% of all respondents (Fig 4). In addition, male survey participants were least likely to attend mental health events (P<0.042, Fisher’s Exact). We chose to conduct these analyses not to further “genderize” our group or its events, but rather to understand what attracts certain subpopulations on campus to different functions and how we can better include all STEMinists (STEM-feminists, regardless of gender) in the future.

Since the collection of this data in 2018, our group has taken steps to address this finding. Specifically, we have established a Faculty Advisory Board (FAB) to help guide the student executive board. The FAB includes a male principle investigator who actively contributes rigorous gender disparity literature to peer review journals. We believe his involvement may encourage other male community members to be more involved. We have additionally taken the step of adding “all genders welcome” on all event advertisements (i.e. campus flyers, emails, and social media broadcasts).

We are encouraged by the fact that male individuals were nominated for two WiSE-hosted campaigns. The first was the 2019 WiSE Mentorship Awards, which aims to recognize invaluable colleagues at the faculty/administrator level and the graduate student/post-doc level who have served as personal or professional mentors to women at CSHL. Historically, despite our efforts, only female nominations were submitted. We are pleased that our community appears to better understand the role of male mentorship for female trainees. Secondly, we have had male nominations for our 2019-2020 election cycle (currently on-going). We are encouraged that this indicates male members of the community are becoming more involved with our events and our leadership.

We further encourage collaboration with diversity specific advocacy groups to address issues that are common for racial and gender minorities within STEM i.e. intersectionality. For example, we have organized an Allyship workshop with CSHL’s Diversity Initiative for the Advancement in STEM (DIAS). Commonalities between this group and our own allow us to build stronger coalitions that can work together toward larger goals. Understanding how to cater to the needs of each group is fundamental for garnering support from the diverse populations that make up any modern academic environment.

### Cultivate mentorship

Mentorship is fundamental for the development and retention of female students in STEM.^12,13^ Identifying strong faculty mentors can help WiSE groups achieve better communication between the group and the administration. Our WiSE Faculty Advisory Board consists of active CSHL research faculty who have demonstrated their commitment to the WiSE mission by advocating for the inception of the group, participation and advocacy for gender studies research, and participation in WiSE events. The primary responsibility of the Faculty Board is to provide mentorship and guidance to WiSE members (the full description of roles and responsibility of the Faculty Board can be found on our website: http://cshlwise.org/about/leadership/).

We encourage other groups to make clear the “what” and “how” of mentorship.^14–16^ Our Faculty Board consists of a mixture of male and female scientists at the assistant, associate, and full professor levels. The diversity of this advisory board is an important factor for its success. For example, selection of male mentors enhances the inclusivity of the group to non-female supporters, and reinforces the message that WiSE is open to all members of the community.

That said, we remind fellow grassroots organizations that faculty members are there to guide and assist but not to lead the student groups. The responsibility of leadership and decision-making should rest in the hands of student leaders. This allows trainees to gain leadership experience while keeping the focus of our organization on issues most strongly affecting our members.

### Reach out beyond your scientific community

The ability to see role models in science from a diverse array of backgrounds, especially at an early age, can impact implicit biases widely. While presenting female role models for younger generations has positive impact,^17^ after several years of doing so we have realized the need to expand these efforts to educate parents, teachers and community leaders, and to develop community contacts/connections for future events. We encourage groups like our own to assess their areas of expertise and consider creative ways to engage their local communities beyond their academic campuses. For example, to include older populations in our education initiatives, we have invited students to bring their parents to our lectures focused on genetically modified organisms (GMOs) with a question-and-answer panel of plant biologists. Such efforts reflect well on the home institution and may provide an important source of independent funding for future initiatives (Fig 6).

**Figure 6.**
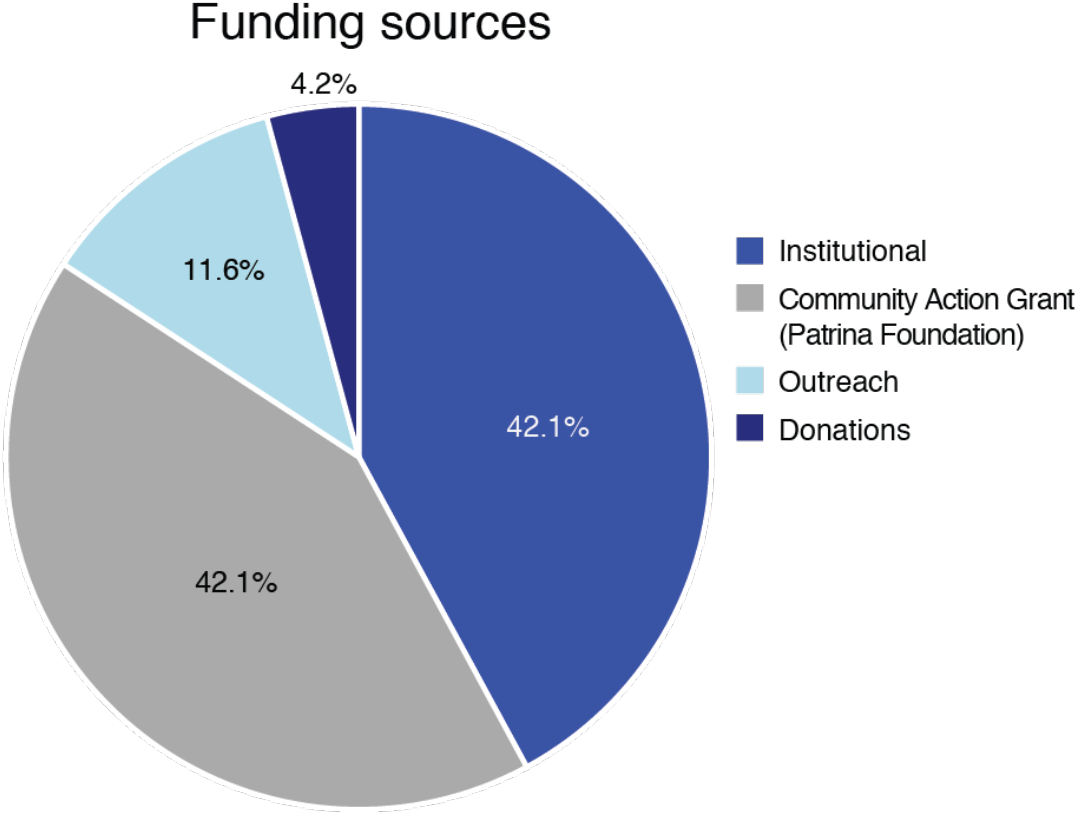
Financial sources of support. Here we have categorized our financial sources of support, namely, funding for specific institutional initiatives, funding from WiSE-hosted outreach events (i.e. instructional education for children or other community members), grant funding (to date, from non-for-profit groups as opposed to government agencies), and donations. We encourage advocacy groups to diversify their funding sources as a means of providing long-term assurance and as a way of promoting institutional support for the group’s efforts.

### Assess, analyze, adapt

In an effort to strengthen our own group, we have gathered, analyzed and discussed the survey data that led to the conclusions drawn here. We hope doing so will serve as an example for other groups to apply their critical, scientific nature towards their passion projects. Whenever possible, we encourage groups to send out follow-up surveys to assess the strengths and weaknesses of specific events; use the **SMART** (specific, measurable, achievable, relevant, and time-based) goal technique to ensure follow-through. Discuss as a team the cost-benefit analysis of hosting each event to order to determine where limited time, effort, and resources are best spent.

### Identify your external resources

There are a growing number of resources to aid grassroots organizations like our own. We encourage groups to familiarize their leadership and members with the currently available online resources from government institutes, academic centers, and private organizations. These tools range from concepts for workshops to data-based reports of gender bias, to funding opportunities. For example, we have found that applications to small community-based initiative grants may be more approachable for grass-roots organizations like our own as opposed to larger national-based opportunities. We have had success in such smaller applications (Fig 6) but caution other groups that all grant applications are competitive and time-demanding, especially when diversity advocacy is a “passion” project undertaken along-side full-time scientific demands.

We provide a more extensive, centralized list of available resources on our website (http://cshlwise.org/resources) but recommend the following as examples: Harvard’s Implicit Bias Training; “The Sexual Harassment of Women: Climate, Culture, and Consequences in Academic sciences, Engineering, and Medicine” published by The National Academies of Sciences, and resources offered on the HeForShe website (https://www.heforshe.org/en).^5,18^

In addition, we recommend following and interacting with similar groups via social media (e.g. Academic Twitter) for example @MeTooSTEM, @500womenscientst, etc. Our own group can be found at @CSHL_WISE.

### Use criticism constructively

We acknowledge the great amount of time and effort required to successfully organize a WiSE group. Facing criticism in the face of this effort is challenging, but nevertheless valuable. Anonymous feedback including critiques from those who generally support the goals of the group should be expected. In addition to this, we want other advocacy leaders to be prepared to face criticism from those who may not currently share the goals of the group, but who are still part of our community. For example

> “WiSE has created a perception that female graduate students do not want to work and they… blame men when they do not have data or [are] not productive in the lab.”

We use this as an opportunity to demonstrate our last point: above all else, persist. Encouraging others to embrace diversity and inclusivity in science takes time and work, but is an important part of improving science as a field. Using critiques like the one above to inspire rather than discourage our work, we continue to modify and amend our group’s focus and events in a continual effort to improve our service and to support the women on our academic campus.

## Conclusion

Here we have shared lessons learned while growing a grassroots organization, targeting specific inequities while balancing inclusivity, and accepting constructive criticism. Our recommendations are by no means a one-size-fits-all model. We believe many of the take-aways supported by our sample data can be generalized to other academic communities. However, we hope that advocacy groups at all stages are encouraged to conduct similar data-driven evaluations of their efforts and can take advantage of our findings at their home institutes so that the STEMinist community at large works resourcefully towards the greater goal of achieving gender parity in STEM.

## Supporting information

Supplementary File 1: The WiSE Survey

Supplementary File 2: WiSE Survey Responses

## Supplemental Materials

Supplemental File 1: The WiSE Survey.

Supplemental File 2: WiSE Survey Responses.

## End Matter

### Author Contributions and Notes

DDR designed the study and collected responses. DDR and ACN manually cleaned data and reviewed prior literature. OHT performed the statistical calculations. MGH advised on data interpretation. DDR drafted the original manuscript; all authors contributed to the writing, editing, finalization of the manuscript, and figure design.

The authors declare no conflicts of interest.

## Acknowledgments

We thank the anonymous participants of our survey at Cold Spring Harbor Laboratory (CSHL). We respectfully acknowledge the founders of the CSHL Women in Science and Engineering (WiSE) organization: Dr. Jaqueline Giovanniello, Dr. Lital Chartarifsky-Lynn, Alexandra Ambrico, and Dr. Leemor Joshua-Tor & Dr. Anne Churchland. We thank Drs. Anne Churchland and Jason Sheltzer for their feedback and comments on this manuscript. Finally, we thank all past and current CSHL WiSE student board members: Brianna Bibel, Tzvia Pinkhasov, Cassidy Danyko, Lyndsey Aguirre, Dr. Grinu Mathew, Dr. Sarah Starosta, and Kaitlin Watrud.

## Notes

#### Summary of Updates

Addition of survey response data to the supplementary files.

