## Supplementary File 1: The WiSE Survey for "Methods for Running a Successful Women-in-STEM Organization on an Academic Campus"

WiSE Needs Survey 2018:

1. What is your current position?

Technician

Graduate student

Post doc

Faculty

Staff

Other (please specify)

2. How long have been at CSHL?

<6 months

6 months-1 year

>1 year

>2 years

>3 years

>4 years

5+ years

3. What is your gender?

Male

Female

Other (please specify)

4. Are you a current member of our group (WiSE)?

Yes

No

5. If you are not a current member, were you previously a member?

Yes

No

N/A- I am currently a member

6. If you are not a current member, tell us why:

N/A- I am a current member

I was previously a member and decided the group wasn't for me…

I was never a member but don't think the group applies to me

I've thought about getting involved but I don't know how

I have too many demands on my time

I don't think the group is productive

I think the group is too political in nature

I don't think it is necessary to have a women-focused group on campus

I don't think I could be helpful for the group

Other (please specify)

7. 7. What do you understand to be the goal(s) of the WiSE group? (check all that apply)

WiSE is a female-only group that promotes women and girls in science and science-related fields

WiSE is a political group aimed at addressing societal and policy issues that lead to gender inequality

WiSE is an open group for everyone at CSHL that promotes women and girls in science and science-related fields

8. On a scale of 1-7 how successful to feel WiSE in reaching your understanding of its goal?
9. How do you think WiSE can improve?

Make the group more inclusive

Improve communication between the group and the CSHL campus

Organize more events

Organize different types of events than those currently hosted

Other (please specify)

10. Which WiSE events have you attended in the past?

Open-mic/mental health night

WiSE summer party/BBQ

WiSE open board meeting/interest meetings

McClintock lectures

Workshops e.g. public speaking, financial etc

Other (please specify)

11. If you have further comments on how we can address the needs of the CSHL community please let us know below!
